# *Phasis*: a software tool for register-resolved discovery of plant phased small RNA loci

**DOI:** 10.64898/2026.07.11.737977

**Authors:** Thales Henrique Cherubino Ribeiro, Atul Kakrana, Vinicius Andrade Maia, Scott Lewis, Blake C. Meyers

## Abstract

Plant *PHAS* locus discovery remains challenging because phasiRNA-producing loci must be distinguished from other sRNA-producing regions with high abundance or apparent periodicity. This problem is especially acute for reproductive 24-*PHAS* loci, which occur within genomes that also produce abundant 24-nt siRNAs from non-*PHAS* regions. We present *Phasis*, an open-source Python software tool for plant *PHAS*-locus discovery from small RNA sequencing data. *Phasis* combines statistical evidence for phased accumulation with locus-level features and a Register-Resolved Locus Interpretation Layer that evaluates whether candidate loci show coherent phased architecture. Across diverse plant datasets, *Phasis* recovered validated or annotated 21- and 24-*PHAS* loci with a strong balance between call-level precision and reference-locus recall, and generally outperformed *PhaseTank* and *ShortStack* in matched benchmark analyses. The register-resolved interpretation layer reduced unsupported calls by separating coherent phased loci from ambiguous sRNA-producing regions. In maize *dcl5* mutant libraries, Phasis showed strong depletion of 24-*PHAS* recovery, supporting DCL5-dependent recovery of reproductive 24-*PHAS* signal. Together, these results support *Phasis* as a biologically interpretable tool for large-scale discovery of plant DCL-dependent phasiRNA loci.

## Introduction

Phased small interfering RNAs (phasiRNAs) are secondary small RNAs (sRNAs) produced from precursor transcripts after a trigger cleavage event establishes a phased register. They are termed secondary because their biogenesis is triggered by an upstream miRNA-guided cleavage event rather than by direct processing of the original precursor alone. In plants, this process depends on conversion of the target RNA into double-stranded RNA and subsequent Dicer-like (DCL) processing, which generates regularly spaced 21- or 24-nt phasiRNAs (Allen et al. 2005; Johnson et al. 2009; Zhan and Meyers 2023). Here, “phased” refers to the register-dependent production of sRNAs at regular intervals from an initiating cleavage site through successive DCL processing, distinguishing phasiRNAs from other DCL products, such as miRNAs, that are processed by DCL enzymes but do not define phased loci. The genomic loci whose transcripts give rise to phasiRNAs are referred to as *PHAS* loci.

Plant *PHAS* loci are diverse in precursor type, developmental context, and evolutionary distribution. Classic *Arabidopsis trans*-acting small interfering RNA (tasiRNA) loci, commonly referred to as *TAS* loci, established the mechanistic link between miRNA-directed cleavage and phasiRNA production (Allen et al. 2005; Peragine et al. 2004; Vazquez et al. 2004) whereas later studies identified phasiRNAs from protein-coding transcripts, including resistance-gene and *PPR* families, as well as abundant reproductive 21- and 24-nt phasiRNAs in many angiosperms (Fei et al. 2013; Xia et al. 2019; Zhai et al. 2011, 2015). Because *PHAS* loci arise from different precursor types and operate in different biological contexts, their discovery can reveal regulatory pathways associated with development, reproduction, and defense. Accurate annotation of these loci is also increasingly important for plant genome resources, which need to represent functional noncoding RNA loci alongside protein-coding genes. The Kronos wheat functional genomic resource provides one example, with genome-wide annotations of both microRNAs and phasiRNAs (Seong et al. 2026).

Computational *PHAS*-locus discovery is challenging because a locus may appear to produce many sRNAs in a sequencing dataset for reasons unrelated to phasiRNA biogenesis. Genuine *PHAS* loci must therefore be distinguished from other genomic regions or transcripts that accumulate sRNA reads but lack coherent DCL-dependent phased processing. A well-supported *PHAS* locus requires more than local and periodic sRNA abundance patterns: the reads should show the expected 21- or 24-nt size class, a coherent phased register, and support across multiple phased positions. We use “phasiRNA signal” to describe this coordinated local pattern, because abundant background sRNA populations, especially heterochromatic 24-nt siRNA loci, can show high abundance or apparent periodicity without representing genuine phasiRNA biogenesis (Polydore et al. 2018). These features favor approaches that combine statistical evidence for phased accumulation with locus-level properties rather than relying on a single score or fixed parameter alone.

Phased sRNA patterns also occur outside plants, most notably in animal piRNA pathways, but these systems differ mechanistically from plant DCL-dependent phasiRNA biogenesis (Anleu Gil and Meyers 2026). Drosophila phased piRNAs are generated through Zucchini-dependent processing (Mohn et al. 2015; Han et al. 2015) and have broader length distributions than plant DCL-dependent phasiRNAs (Anleu Gil and Meyers 2026). These differences provide a useful boundary for evaluating whether a plant-focused *PHAS*-identification tool captures DCL-dependent phasiRNA architecture rather than generic or apparent periodic sRNA accumulation.

A set of command-line tools has been developed for de novo *PHAS*-locus discovery, including *PhaseTank*, *PhasiHunter*, *DIGITAL*, and earlier versions of *ShortStack* (Guo et al. 2015; Bu et al. 2023; Feng et al. 2023; Axtell 2013). However, robust inference across heterogeneous plant datasets remains difficult because *PHAS* classes differ in abundance, complexity, strand bias, phase length, and background sRNA composition. This creates a need for software that is reproducible across datasets while still allowing evidence thresholds and interpretation to reflect biological context. We previously developed a computational tool called *PHASIS* for de novo discovery and characterization of phasiRNA-generating loci (Kakrana et al. 2017). Here, we describe *Phasis* version 2, hereafter *Phasis*, a substantially redesigned and expanded successor focused on robust plant *PHAS*-locus discovery and interpretation from sRNA sequencing data.

*Phasis* identifies candidate *PHAS* loci by evaluating whether sRNAs associated with a genomic region show features expected from DCL-dependent phased processing, including enrichment for a dominant sRNA size class, regularly spaced reads, sufficient abundance, and evidence that the signal is unlikely to arise from random sRNA accumulation. Candidate loci are then evaluated by a Register-Resolved Locus Interpretation Layer (RRL). Here, “register-resolved” means that *Phasis* does not treat a candidate locus only as a broad interval with a single score; instead, it inspects phased positions within the locus to identify the dominant register, define the main phased unit, and distinguish coherent phased structure from secondary, overlapping, or unsupported signals. Using benchmark datasets across multiple plant systems, we show that *Phasis* recovers validated 21- and 24-*PHAS* loci with strong false-positive control, performs well relative to existing tools, and reports biologically interpretable locus structures. We further use maize *dcl5* mutant libraries to validate pathway-specific 24-*PHAS* recovery and Drosophila piRNA-producing libraries as a boundary test for the plant-focused scope of the software.

## Results

### Overview of the Phasis Software Workflow

We developed *Phasis* to identify plant *PHAS* loci from sRNA sequencing (sRNA-seq) data by evaluating whether sRNAs originating within genomic or transcript-derived clusters show the size, abundance, and positional organization expected from DCL-dependent phased processing. The workflow has two stages (Fig. 1A). First, *Phasis* performs Initial *PHAS* Candidate Detection, in which sRNA-producing clusters are identified and scored using computed evidence features such as size-class enrichment, local abundance, strand bias, and phased-register support. Second, candidate loci are examined by a Register-Resolved Locus Interpretation Layer (RRL), which summarizes how the phasiRNA signal is organized within each candidate locus. Library-resolved summaries of *PHAS* detection and abundance can then be used to compare phased loci across samples or biological conditions, providing a genome-wide view of candidate *PHAS* signal across libraries (Fig. 1B).

**Figure 1.**
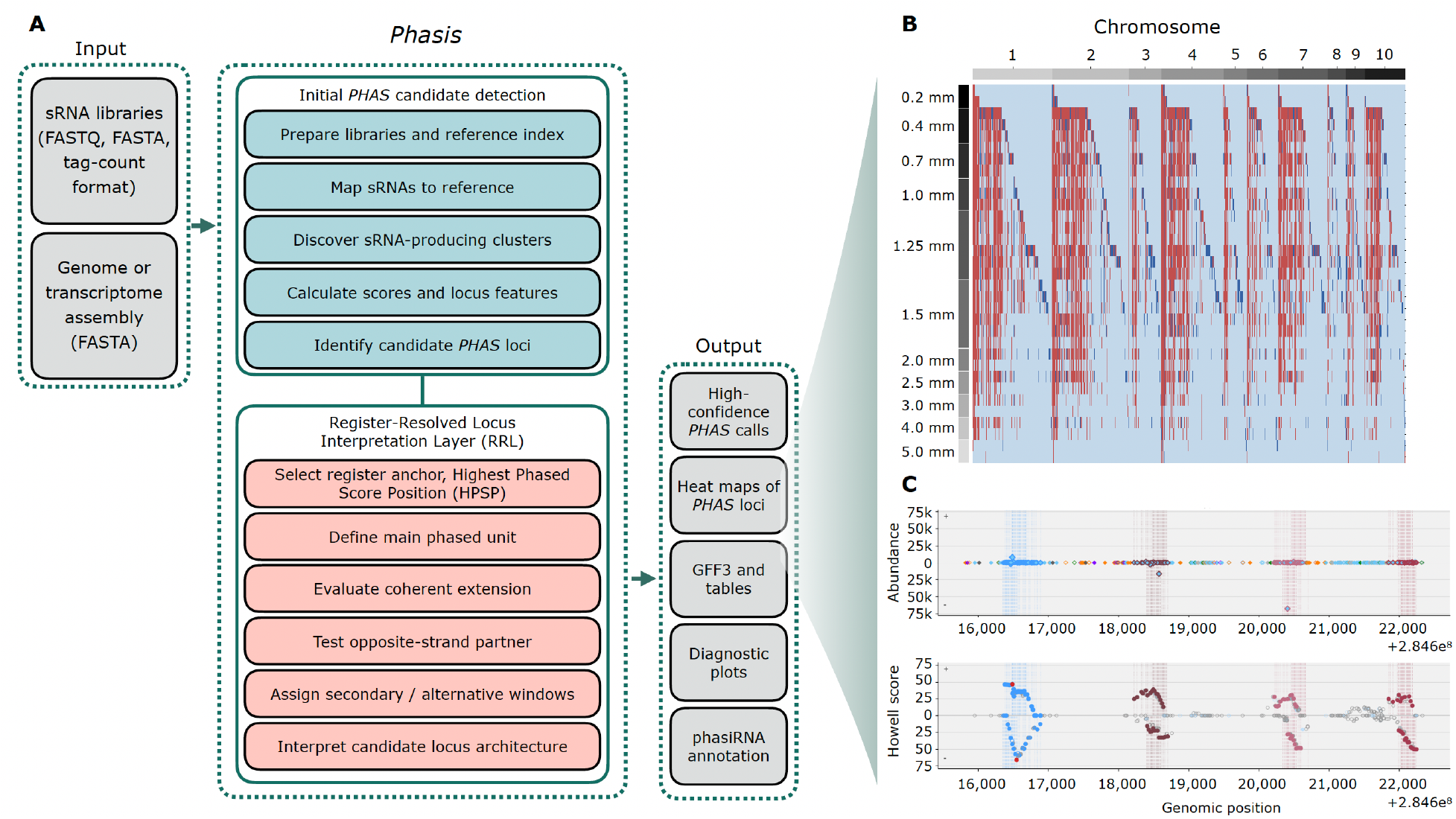
*Phasis* workflow and representative register-resolved outputs. **(A)** *Phasis* analyzes sRNA libraries together with a genome or transcriptome reference to identify candidate *PHAS* loci. After library preparation, reference indexing, sRNA mapping, cluster discovery, and locus-level evidence calculation, initial candidate *PHAS* loci are passed to the Register-Resolved Locus Interpretation Layer (RRL). RRL selects a register anchor based on the highest phased-score position (HPSP), defines the main phased unit, evaluates coherent register extension, tests opposite-strand partners, assigns secondary or alternative windows, and interprets candidate locus architecture. Outputs include high-confidence *PHAS* calls, *PHAS* locus heat maps, GFF3 (a standard genome annotation format that describes genomic features) and tabular files, diagnostic plots, and phasiRNA annotations. **(B)** Representative genome-wide *Phasis* heat map showing candidate *PHAS* signal across chromosomes and sRNA libraries. Rows correspond to developmental-stage libraries, columns correspond to genomic positions ordered by chromosome, and colored marks indicate sRNA-producing intervals summarized across genomic position. Red indicates candidate *PHAS* loci, dark blue indicates sRNA-producing loci not called as *PHAS*, and light blue indicates weak or no detected signal. **(C)** Representative locus-level diagnostic output. The top plot shows local sRNA abundance across genomic position, with reads displayed above and below the midline according to mapped strand. This strand-separated display reflects the double-stranded precursor context expected for plant phasiRNA biogenesis and helps distinguish coherent phased signal from unsupported accumulation. Colors mark sRNA size classes and shaded vertical bands mark register-resolved phased windows. The bottom plot shows Howell-score support across the same genomic interval, again separated by strand, illustrating where phased support is concentrated relative to the interpreted register-resolved windows. A more detailed explanation of the color and symbol conventions used in these diagnostic plots, together with representative high-confidence, ambiguous, and unsupported candidate loci, is provided in Fig. S1.

During Initial *PHAS* Candidate Detection, *Phasis* evaluates evidence for phasiRNA accumulation using multiple locus-level features. The *Phasis* score summarizes statistical support for phased accumulation, whereas the Howell score provides an adapted register-level phased-score metric based on the formula described by Howell et al. (2007). In *Phasis*, the Howell score allows strand-specific positional variance of plus or minus one nucleotide at the 5′ end of the expected phase register, an implementation choice intended to tolerate limited positional variation while preserving register-level evidence for DCL-dependent phased processing. Detailed definitions of these features and the full *Phasis* score calculation are provided in the Supplemental Methods.

The RRL adds a post-detection interpretation step. Rather than treating each candidate as a single undifferentiated interval, this layer examines the local phased pattern and summarizes whether the signal forms a clear main phased unit, includes opposite-strand support, or contains additional supported phased windows. *Phasis* also reports locus-level diagnostic plots, inspired by the web-based MPSS/Plant small RNA database phasing-analysis viewer described by Nakano et al. (2020), that display local sRNA abundance, strand organization, size-class composition, register-resolved windows, and Howell-score support across the candidate interval (Fig. 1C). A detailed guide to these diagnostic plots, including color and symbol conventions and representative high-confidence, ambiguous, and unsupported candidate loci, is provided in Fig. S1. Together, these outputs allow *Phasis* to report both candidate-level evidence and a biologically interpretable model of phasiRNA organization within each locus.

### Phasis *Improves the Precision–Recall Balance Across Plant* PHAS *Benchmarks*

We first evaluated whether *Phasis* could recover validated or annotated *PHAS* loci across diverse plant datasets. The benchmark set included maize and coffee 21-*PHAS* and 24-*PHAS* references, curated reproductive 24-*PHAS* references from petunia and columbine, and annotated Arabidopsis *TAS* loci. These datasets were selected to represent multiple discovery contexts, including reproductive 24-nt phasiRNA-producing loci, protein-coding or noncoding phased precursors, and a negative 24-*PHAS* context in *Arabidopsis*, which lacks the miR2275–DCL5 reproductive 24-nt phasiRNA pathway. This range of datasets also tested performance in 24-nt-rich plant backgrounds, where abundant 24-nt siRNA populations can lead to false-positive 24-*PHAS* annotation (Polydore et al. 2018). Here, individual-library evidence refers to loci detected in one or more constituent libraries, whereas pooled-library evidence refers to loci detected after libraries were combined before candidate detection.

Across these plant benchmarks, *Phasis* recovered expected *PHAS* loci with a strong balance between reference recovery and false-positive control. In petunia flower bud libraries (Xia et al. 2019) aligned to the reference genome described by Bombarely et al. (2016), pooled-library *Phasis* analysis recovered 367 of 440 curated 24-*PHAS* reference loci with 11 false-positive calls, corresponding to a precision of 0.97, recall of 0.83, and F1 score of 0.90. The F1 score summarizes the balance between precision and recall as their harmonic mean. These results support robust 24-*PHAS* identification despite the alternative phased-register architecture previously described in petunia (Xia et al. 2019). In columbine, using the genome assembly described by Filiault et al. (2018), *Phasis* recovered 466 of 576 curated 24-*PHAS* reference loci in pooled-library analysis, with a precision of 0.92, recall of 0.81, and F1 score of 0.87.

Arabidopsis provided a complementary validation context. Rather than benchmarking reproductive *PHAS* recovery against a positive reference, we tested whether *Phasis* recovered well-characterized *TAS*-associated 21-*PHAS* loci while avoiding unsupported interpretation of 24-*PHAS* candidates in a species lacking the miR2275–DCL5 reproductive 24-nt phasiRNA pathway. In sRNA libraries from 12-day-old Arabidopsis seedlings (Li et al. 2016), pooled-library analysis recovered six of eight annotated *TAS* loci. *AtTAS3b* and *AtTAS3c* were not recovered by either evidence mode, possibly reflecting limited local phasiRNA abundance in bulk seedling libraries given the spatially restricted activity of the miR390–*TAS3* pathway during lateral-root development (Marin et al. 2010). *Phasis* also produced no high-confidence individual-library 24-*PHAS* calls in Arabidopsis, consistent with the absence of the miR2275–DCL5 reproductive 24-nt phasiRNA pathway in this species.

We next compared pooled-library *Phasis* outputs with ShortStack v3.8.5 and PhaseTank v1.0 using matched benchmark datasets and the same curated reference sets (Fig. 2A–C; Supplemental Table S1). These comparisons were restricted to combined-evidence outputs because the comparator tools do not provide the same individual-library evidence mode as *Phasis*. Across these matched analyses, *Phasis* generally provided the strongest balance between reference recovery and false-positive control, as summarized by F1 score (Fig. 2A), while the true-positive and false-positive call counts are shown in Fig. 2B. In petunia 24-*PHAS* analysis, *Phasis* achieved an F1 score of 0.90, compared with 0.64 for *ShortStack* and 0.58 for *PhaseTank*. In columbine 24-*PHAS* analysis, *Phasis* again had the highest F1 score, 0.87, compared with 0.41 for *ShortStack* and 0.70 for *PhaseTank*.

**Figure 2.**
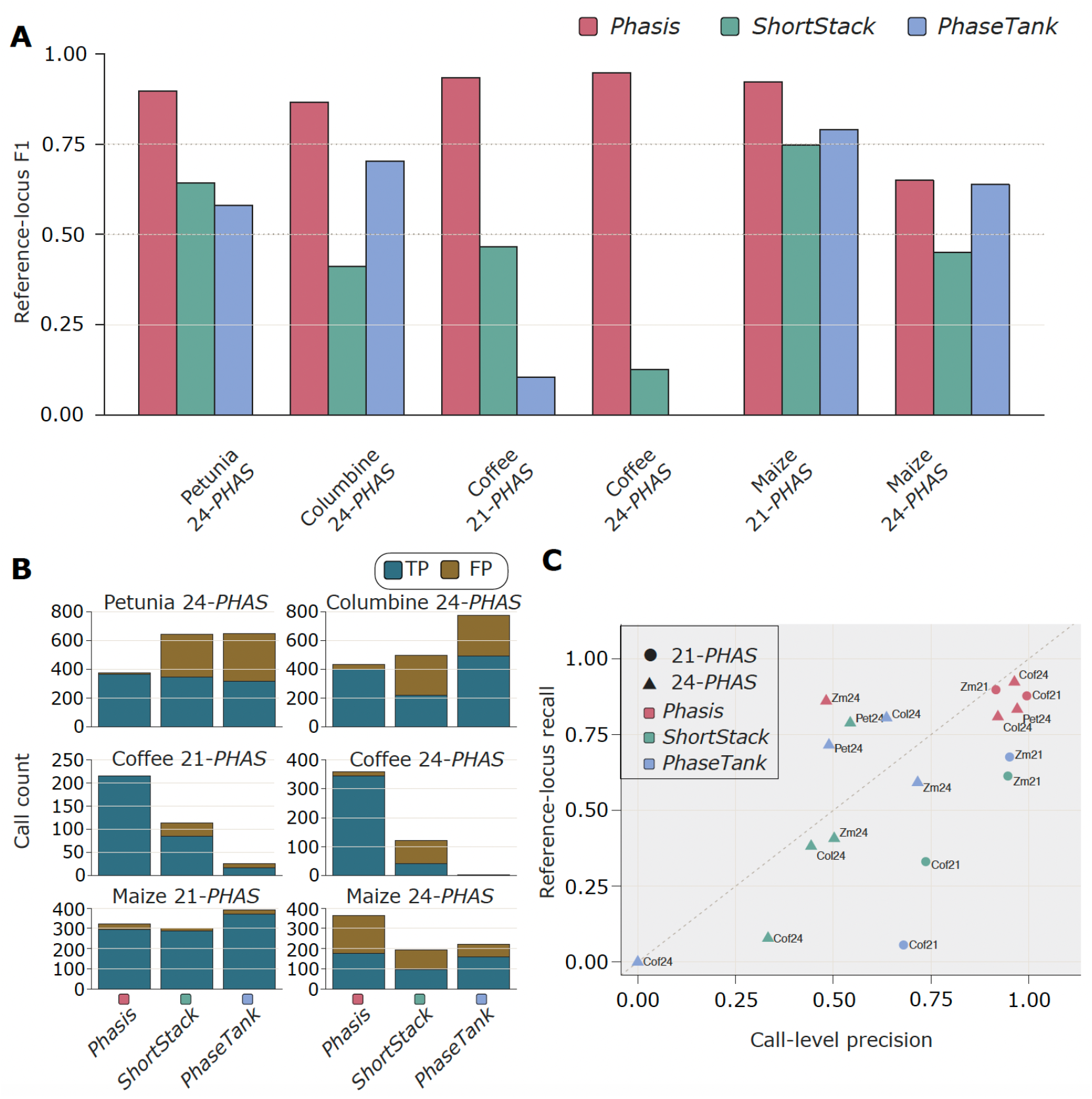
Pooled-library benchmark comparison of *Phasis*, *ShortStack*, and *PhaseTank* across plant *PHAS* datasets. **(A)** F1 scores calculated from call-level precision and reference-locus recall after *PhasMatch* comparison against curated reference sets for petunia 24-*PHAS*, columbine 24-*PHAS*, coffee 21-*PHAS* and 24-*PHAS*, and maize wild-type anthers 21-*PHAS* and 24-*PHAS* benchmarks. To enable direct comparison, this benchmark shows only pooled-library *Phasis* results, because ShortStack and PhaseTank were evaluated using pooled-library input and do not provide the individual-library evidence mode used by *Phasis*. Software colors are consistent across all panels and figures. **(B)** Threshold-positive call counts for each pooled-library benchmark and tool, split into true-positive and false-positive calls. True positives are calls matched to curated reference loci, whereas false positives are threshold-positive calls not matched to the corresponding reference set. **(C)** Call-level precision versus reference-locus recall for each pooled-library benchmark and tool. Circles indicate 21-*PHAS* analyses and triangles indicate 24-*PHAS* analyses. Point colors indicate software as in panel A and are used consistently across figures. The dashed diagonal marks equal precision and recall; points closer to the upper-right indicate stronger joint call-level precision and reference-locus recovery.

The strongest performance differences were observed in coffee. For coffee 21-*PHAS* loci, *Phasis* achieved an F1 score of 0.93, compared with 0.47 for *ShortStack* and 0.10 for *PhaseTank*. For coffee 24-*PHAS* loci, *Phasis* achieved an F1 score of 0.95, whereas *ShortStack* and *PhaseTank* had F1 scores of 0.13 and 0, respectively. In maize 21-*PHAS* analysis, Phasis also had the highest F1 score, 0.92, followed by *PhaseTank* at 0.79 and *ShortStack* at 0.75. Maize 24-*PHAS* remained the main exception in the pooled-library comparator benchmark, where *Phasis* and *PhaseTank* had similar F1 scores, 0.65 and 0.64, respectively, while *ShortStack* was lower at 0.45.

The precision–recall relationship was consistent with these benchmark results: across datasets, *Phasis* generally showed high precision and high recall, whereas comparator outputs more often showed reduced recall or lower precision depending on the dataset (Fig. 2C). Together, these plant benchmarks show that *Phasis* generally improved the balance between reference recovery and false-positive control across diverse *PHAS* discovery contexts. The maize 24-*PHAS* comparator result also indicates that pooled-library benchmarking captures only one operating mode of *Phasis*. We therefore separately evaluated how evidence mode affects recovery and precision.

### *Evidence Mode Affects* PHAS *Recovery and Precision*

We next evaluated how individual-library and pooled-library evidence affected *PHAS* recovery under the final feature-threshold-enabled and RRL configuration. Across maize and coffee, pooled-library evidence generally increased reference-locus recovery, but the magnitude and precision effect depended strongly on dataset and phase class (Fig. 3A; Supplemental Table S2). In maize, pooled-library evidence modestly increased recall for both 21-*PHAS* loci, from 0.82 to 0.88, and 24-*PHAS* loci, from 0.82 to 0.86. In coffee, the effect was much larger, increasing recall from 0.36 to 0.88 for 21-*PHAS* loci and from 0.09 to 0.92 for 24-*PHAS* loci. These gains were especially clear for coffee 24-*PHAS*, where pooled-library evidence recovered 459 of 497 reference loci with a precision of 0.96 and F1 score of 0.95. By contrast, maize 24-*PHAS* pooled-library analysis increased recall but reduced precision, showing that pooled evidence can improve recovery while altering the precision–recall balance (Fig. 3A; Supplemental Table S2).

**Figure 3.**
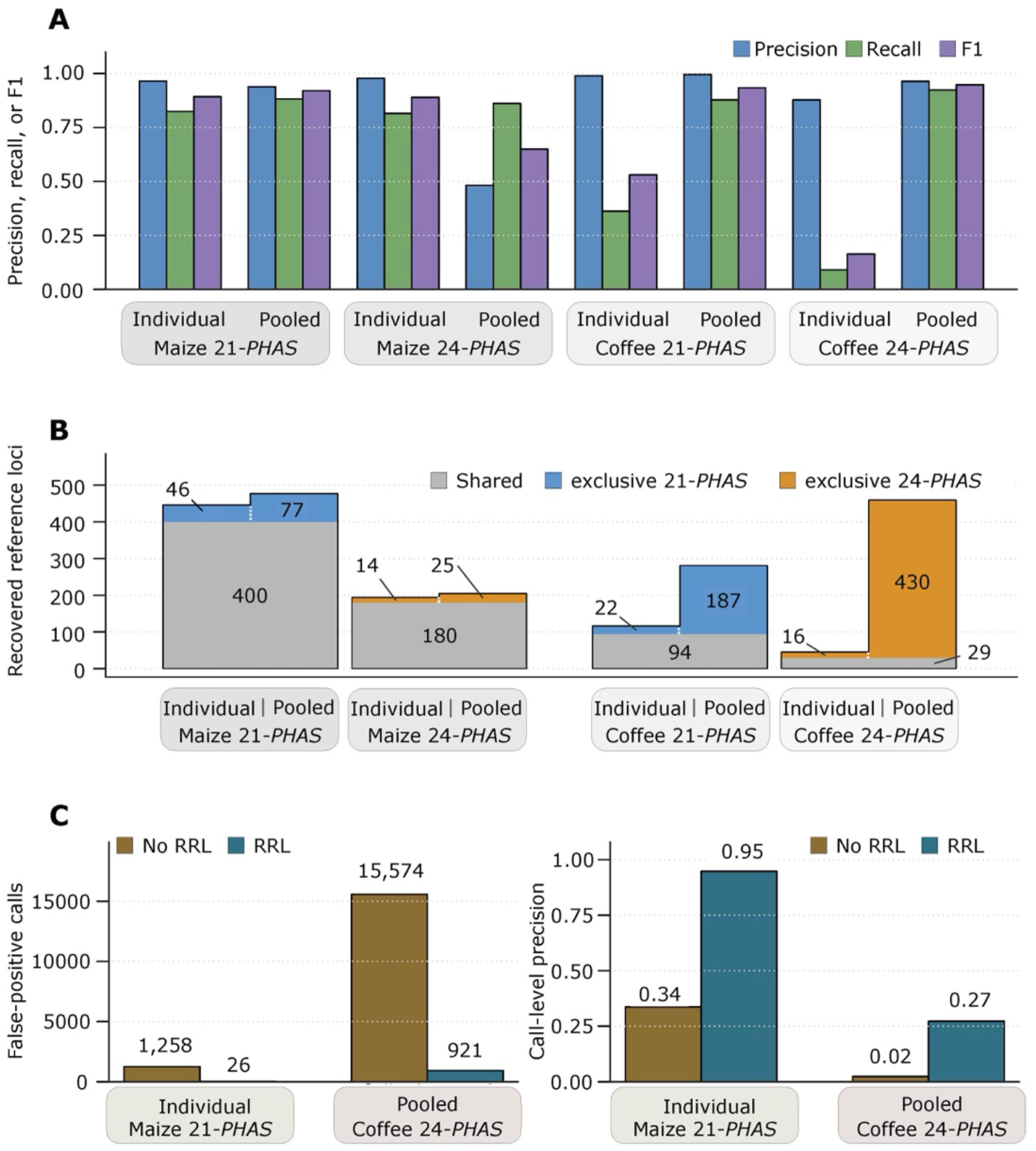
Evidence mode and register-resolved interpretation affect *PHAS* recovery and false-positive control. **(A)** Call-level precision, reference-locus recall, and F1 score for individual-library and pooled-library *Phasis* analyses under the default configuration. Results are shown for maize and coffee 21-*PHAS* and 24-*PHAS* benchmarks. **(B)** Reference-locus overlap between individual-library and pooled-library evidence under the same feature-threshold-enabled and RRL-enabled configuration. Gray segments indicate reference loci recovered by both evidence modes. Colored segments indicate loci recovered only by one evidence mode, with blue denoting 21-*PHAS* benchmarks and gold denoting 24-*PHAS* benchmarks. Numbers indicate recovered reference loci in each segment. Pooled-library evidence increased total recovery in several contexts but was not a strict superset of individual-library detection. **(C)** Effect of RRL on false-positive control under feature-threshold-disabled detection settings. Left, false-positive call counts with and without RRL for maize 21-*PHAS* individual-library analysis and coffee 24-*PHAS* pooled-library analysis. Right, corresponding call-level precision values. RRL reduced false-positive inflation and improved call-level precision by requiring coherent register-resolved phased architecture.

Overlap analysis using the reference loci as the baseline showed that detection from pooled libraries was not a strict superset of detection from individual libraries (Fig. 3B). In maize, pooling libraries recovered more reference loci overall, but analyses of individual libraries still detected 46 21-*PHAS* and 14 24-*PHAS* loci that were missed after pooling. The same pattern was observed in coffee: pooling libraries strongly increased recovery, but analyses of individual libraries still detected 22 21-*PHAS* and 16 24-*PHAS* loci that were not recovered from the pooled data. Thus, pooling changed the evidence profile rather than simply adding statistical power to detection from individual libraries.

The additional plant validation datasets extended this pattern beyond the maize and coffee benchmarks (Supplemental Table S3). In petunia, pooling libraries substantially increased recovery of curated reproductive 24-*PHAS* loci while retaining high precision. In columbine, pooling also increased recovery of reference loci, but it produced a larger number of false positives and reduced precision relative to analyses of individual libraries. In *Arabidopsis*, pooling rescued one additional annotated *TAS* locus, *AtTAS4*, that was not recovered from the individual libraries. These results indicate that the evidence mode should be selected according to the dataset structure and biological question: analyses of individual libraries preserve sample-specific signals, whereas pooling can recover weak or distributed signals but may also change the false-positive profile.

### Register-Resolved Interpretation Controls False-Positive Inflation

Because pooled-library evidence could increase recovery while also altering the false-positive profile, we next tested how much the RRL contributed to false-positive control. Maize and coffee 21-*PHAS* and 24-*PHAS* benchmarks were evaluated under matched configurations that varied feature-threshold status, RRL status, and evidence mode. Feature-threshold-disabled analyses represented a deliberately permissive candidate-detection setting, allowing us to test whether RRL could reduce unsupported calls after broad initial detection. The full benchmark, including all feature-threshold-enabled/disabled, RRL present/absent, individual-library, and pooled-library configurations, is provided in Supplemental Table S2.

Across the full benchmark, RRL had its strongest effect under permissive settings in which feature thresholds were disabled. In maize 21-*PHAS* individual-library analysis, feature-threshold-disabled detection without RRL produced 1,258 false-positive calls and a call-level precision of 0.34, whereas adding RRL reduced false-positive calls to 26 and increased call-level precision to 0.95, with only a small change in reference-locus recall. The same pattern was especially clear in 24-*PHAS* analyses with pooled-library evidence, where the number of unsupported calls was highest. For example, in coffee 24-*PHAS* pooled-library analysis, feature-threshold-disabled detection without RRL produced 15,574 false-positive calls and a call-level precision of 0.02; adding RRL reduced false-positive calls to 921 and increased call-level precision to 0.27 while retaining high reference-locus recall, illustrating the false-positive-control effect summarized in Fig. 3C.

These results show that RRL reduces false-positive inflation when initial candidate detection is intentionally broad. By assessing phased-register consistency rather than relying solely on signal abundance, RRL evaluates whether candidate loci contain coherent register-resolved phased architecture, allowing *Phasis* to separate better-supported *PHAS* calls from unsupported or ambiguous sRNA-producing regions. We next asked whether these high-confidence calls also reflected expected pathway biology in maize *dcl5* mutant libraries.

### dcl5 *Mutant Benchmarking Supports DCL5-Dependent 24-*PHAS *Recovery*

We next used maize *dcl5* mutant anther libraries to test whether *Phasis* recovery of 24-*PHAS* loci reflected the expected DCL5-dependent biology of reproductive 24-nt phasiRNAs. The analysis compared W23 2.0 mm anther libraries with stage-matched homozygous *dcl5* mutant 2.0 mm anther libraries, corresponding to the meiotic window in which 24-nt reproductive phasiRNAs are expected to accumulate (Teng et al. 2020).

Under the fixed-threshold and RRL configuration, homozygous *dcl5* anther libraries showed strong depletion of maize 24-*PHAS* recovery relative to W23 libraries (Fig. 4A). This reduction was observed for both the full maize 24-*PHAS* reference set and the meiotic-enriched 24-*PHAS* reference subset, consistent with the expected dependence of reproductive 24-nt phasiRNA accumulation on DCL5.

**Figure 4.**
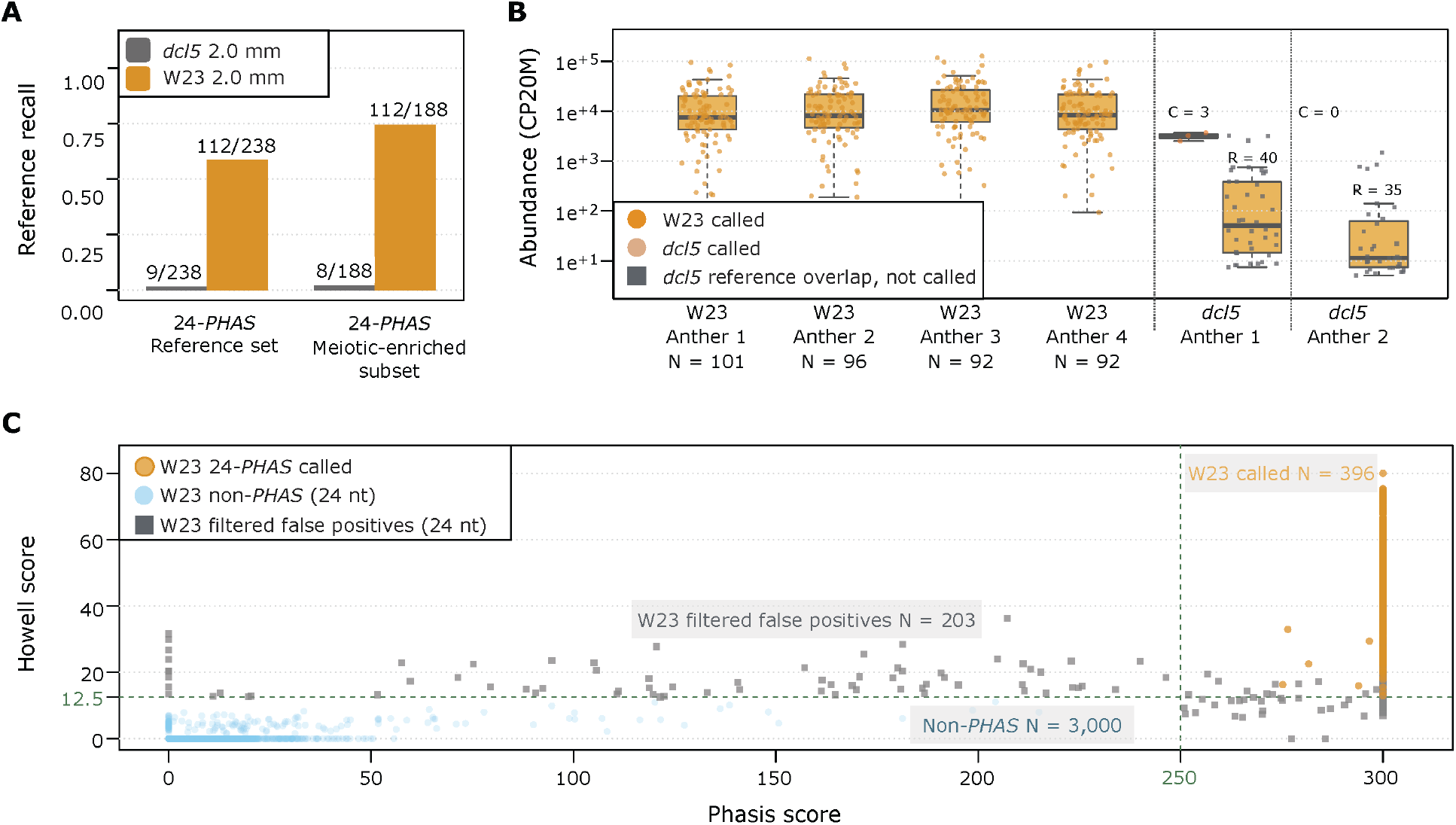
*dcl5* mutant analysis in maize supports pathway-specific recovery of reproductive 24-*PHAS* loci. **(A)** Reference-locus recall in W23 2.0-mm anther libraries and stage-matched homozygous *dcl5* mutant 2.0-mm anther libraries under the default configuration. **(B)** CP20M-normalized abundance distributions for called 24-*PHAS* loci and reference-overlapping non-called loci in W23 and homozygous *dcl5* anther libraries. Raw CP20M values are displayed on a log-scaled y-axis for visualization. W23 called loci are shown in gold, *dcl5* called loci in orange, and *dcl5* reference-overlapping non-called loci as gray points over yellow box plots. N indicates W23 called loci, C indicates *dcl5* called loci, and R indicates *dcl5* reference-overlapping non-called loci. **(C)** *Phasis*-score versus Howell-score diagnostic plot for W23 24-nt candidate filtering. Dashed lines mark the fixed-threshold criteria used for high-confidence 24-*PHAS* calling. W23 called 24-*PHAS* loci are concentrated in the high *Phasis*-score and high Howell-score region, whereas filtered false positives and non-*PHAS* background loci show insufficient or discordant evidence.

The abundance profiles provided a complementary view of this depletion. W23 libraries contained many high-abundance 24-*PHAS* calls, whereas homozygous *dcl5* anther libraries contained few high-confidence 24-*PHAS* calls (Fig. 4B). In the dcl5-1 homozygous 2.0-mm anther library, only three loci were still called as 24-*PHAS* and showed higher median CP20M abundance than reference-overlapping loci that remained detectable but did not pass high-confidence calling criteria. In the *dcl5*-2 homozygous 2.0-mm anther library, no 24-*PHAS* loci passed these criteria. Together, these abundance patterns support strong depletion of the 24-*PHAS* signal in homozygous *dcl5* anthers, with only a small number of high-abundance reference-overlapping loci remaining detectable as high-confidence calls.

Score-space analysis further separated supported 24-*PHAS* calls from threshold-conflict candidates and non-*PHAS* background clusters (Fig. 4C). Validated 24-*PHAS* calls were concentrated in the high *Phasis*-score/high Howell-score region, whereas many filtered candidates showed discordant evidence, including high Howell score with low *Phasis* score or high *Phasis* score with insufficient Howell-score support. Together, the reference-recovery, abundance, and score-space analyses indicate that Phasis detects a DCL5-dependent 24-*PHAS* signal rather than simply calling abundant 24-nt sRNA clusters.

### *Drosophila Scan Defines a Mechanistic Boundary for* Phasis

We applied *Phasis* to sRNA libraries from *Drosophila melanogaster* ovarian tissues, a species and organ known to produce phased piRNAs through Zucchini-dependent processing (Han et al. 2015; Mohn et al. 2015). Because piRNAs span a broader size range than plant DCL-dependent phasiRNAs, we scanned phase periodicities from 23 to 30 nt. Across this scan, *Phasis* reported only three candidates: one 25-nt candidate and two 26-nt candidates.

Manual inspection and coordinate-level annotation based on the FlyBase annotation release 6.54 (Öztürk-Çolak et al. 2024) indicated that these calls did not represent convincing plant-like *PHAS* loci. Their apparent periodicity was supported by limited local signal rather than by distributed register-wide phased support across a coherent locus. One candidate occurred in a repeat-rich region containing a transposable-element insertion and *DNAREP1_DM* Helitron sequence, another contained multiple simple repeats, and the third was dominated by a single 25-nt sequence without broader phased support.

These results define a mechanistic boundary for *Phasis*. Mechanistically distinct sRNA populations, including piRNA-sized or repeat-associated signals, can occasionally produce fixed-spacing patterns that satisfy automated plant-oriented criteria. However, these candidates did not show the architecture expected for plant DCL-dependent *PHAS* loci when evaluated with locus plots and genomic annotations. *Phasis* is therefore intended for plant DCL-dependent phasiRNA discovery rather than for general detection of phased or periodic animal sRNA production.

## Discussion

*Phasis* was designed to support the conservative and biologically interpretable discovery of plant *PHAS* loci from sRNA-seq datasets. The results show that combining Initial *PHAS* Candidate Detection with register-resolved interpretation improves the separation of high-confidence *PHAS* calls from ambiguous or unsupported sRNA-producing regions. Across plant benchmarks, *Phasis* recovered validated 21- and 24-*PHAS* loci with a strong balance between reference recovery and false-positive control, and comparator analyses showed improved performance relative to existing tools in most benchmark contexts.

The benchmark results also show that evidence mode affects *PHAS* recovery. Pooled-library analysis often recovered additional validated loci, especially when phasiRNA signal was weak or distributed across libraries, but it was not a strict replacement for individual-library analysis. Individual-library evidence preserved sample-specific signals and recovered some loci that were missed after pooling. Thus, evidence mode should be selected according to dataset structure and biological question rather than assuming that one approach is always more sensitive than the other.

The RRL and *dcl5* analyses support the importance of evaluating *PHAS* candidates beyond abundance or apparent periodicity alone. RRL reduced false-positive inflation by requiring coherent register-resolved phased architecture, while the maize *dcl5* comparison showed strong depletion of 24-*PHAS* recovery in homozygous mutant anther libraries. Together, these analyses indicate that *Phasis* detects pathway-consistent 24-*PHAS* signal.

A practical consideration is that conservative, register-resolved analysis can require substantial computational resources, particularly for samples enriched in 24-nt sRNAs analyzed in 24-*PHAS* mode. As a general guideline, we recommend allocating approximately 4 GB of RAM per library, although highly abundant or deeply mapped datasets may require more. In some 24-nt-enriched datasets, memory use in intermediate stages approached approximately 1.14 GiB per million mapped reads, and users should therefore adjust memory allocation and expected runtimes according to library size, mapping depth, and phase mode.

The *Drosophila* scan clarifies the intended scope of the software. Mechanistically distinct sRNA populations, including animal piRNAs and repeat-associated signals, can occasionally produce fixed-spacing patterns that satisfy automated plant-oriented criteria. These candidates require biological and annotation-based interpretation and should not be treated as evidence that *Phasis* has the capacity to be a general detector of animal phased sRNA pathways. *Phasis* is therefore best used for plant DCL-dependent phasiRNA discovery, where periodicity can be interpreted together with size class, abundance structure, strand/register support, and known pathway biology.

Together, these results support *Phasis* as a tool for register-resolved discovery and interpretation of plant *PHAS* loci across diverse sRNA-seq datasets. To support manual review and biological interpretation, *Phasis* also generates integrated locus-level diagnostic plots that summarize local sRNA abundance, size-class composition, strand organization, phased-register support, and Howell-score evidence. These visual summaries provide a practical check of candidate architecture and help distinguish coherent *PHAS* loci from overlapping, weakly supported, or ambiguous sRNA-producing regions.

## Materials and Methods

### Small RNA Datasets and Reference Genomes

Small RNA sequencing datasets were selected to represent multiple plant *PHAS* discovery contexts and one non-plant boundary case. Maize and coffee datasets were used for the main 21-*PHAS* and 24-*PHAS* benchmarks (Zhai et al. 2015; Cherubino Ribeiro et al. 2024); petunia and columbine were used as additional reproductive 24-*PHAS* validation datasets (Xia et al. 2019); Arabidopsis seedlings were used for annotated *TAS*-associated 21-*PHAS* recovery and as a negative 24-*PHAS* context (Li et al. 2016); and Drosophila ovarian libraries were used as a mechanistically distinct piRNA-producing system (Han et al. 2015).

For each species, sRNA reads were analyzed against the genome assembly used for the corresponding reference set and benchmark. Arabidopsis *TAS* validation was performed using Araport11 *TAS* gene annotations. Dataset accessions and sample metadata are provided in Supplemental Table S4. Curated *PHAS* locus annotations and coordinates used for validation are provided in Supplemental Table S5. Representative analysis workflows are provided in Supplemental File S1.

### Phasis *Analyses and Evidence Modes*

*Phasis* analyses were performed with phase length set according to the *PHAS* class or small-RNA pathway being evaluated. Plant analyses focused on 21-nt and 24-nt phase lengths, corresponding to the plant phasiRNA classes analyzed in this study. Drosophila analyses used a phase-length scan from 23 to 30 nt to encompass the broader size range of ovarian piRNAs. Unless otherwise indicated, *Phasis* calls discussed as high-confidence *PHAS* loci refer to outputs passing the fixed-threshold and register-resolved interpretation criteria used for the corresponding benchmark.

Two evidence modes were evaluated where appropriate. In individual-library mode, *PHAS* calling was performed separately for each sRNA library, preserving library-specific phasiRNA accumulation, abundance, and register structure. In pooled-library mode, libraries belonging to the same dataset were analyzed as combined evidence before candidate detection, allowing weak signals distributed across libraries to contribute to the same locus. These modes were interpreted as complementary rather than hierarchical alternatives. When both evidence modes were available, overlap between recovered reference loci was evaluated to determine whether pooled-library detection acted as a strict superset of individual-library detection or provided a distinct evidence profile.

### *Reference Sets and* PhasMatch *Benchmarking*

Species-specific reference coordinates were used to evaluate *Phasis* performance for each benchmarked species and phase class. These included curated maize, coffee, petunia, and columbine *PHAS* reference sets and Araport11 *TAS* gene coordinates for Arabidopsis. Arabidopsis 24-*PHAS* outputs were interpreted as a negative biological context because this species lacks the miR2275–DCL5 reproductive 24-nt phasiRNA pathway.

*Phasis* calls were compared with reference loci using the *Phasis* supporting tool *PhasMatch*. Each *PHAS* call was treated as a coordinate-defined call. Calls that overlapped a curated reference locus were counted as true-positive calls, whereas calls without reference support were counted as false-positive calls. Reference loci overlapped by one or more calls were counted as recovered reference loci, whereas reference loci not overlapped by any call were counted as unrecovered reference loci.

These outputs were used to calculate call-level precision, reference-locus recall, and F1 score. Precision was evaluated at the call level, whereas recall was evaluated at the reference-locus level. For analyses comparing individual-library and pooled-library evidence, reference-locus recovery was also summarized as the overlap between reference loci recovered in both evidence modes, recovered only in individual-library analyses, recovered only in pooled-library analyses, or not recovered by either mode. This reference-side overlap was used to determine whether pooled-library evidence behaved as a strict superset of individual-library evidence or provided complementary support.

### dcl5 *Mutant Validation and CP20M Abundance Analysis*

Maize *dcl5* mutant libraries were used to evaluate whether *Phasis* recovery of 24-*PHAS* loci reflected the expected DCL5-dependent biology of reproductive 24-nt phasiRNAs. The analysis compared W23 2.0-mm anther libraries with stage-matched homozygous *dcl5* mutant 2.0-mm anther libraries. W23 and *dcl5* libraries were analyzed using the same fixed-threshold and register-resolved interpretation criteria applied in the maize benchmark analyses.

Reference recovery was evaluated against the full maize 24-*PHAS* reference set and against a meiotic-enriched 24-*PHAS* subset. Maize 21-*PHAS* reference recovery was evaluated in the same *dcl5* libraries as an internal control for general 21-nt phasiRNA detection. Recovery of reference loci was assessed using *PhasMatch*.

For abundance comparisons, 24-*PHAS* calls and reference-overlapping non-called loci were quantified using total locus abundance reported by *Phasis*. Abundance was normalized as CP20M by dividing total locus abundance by the corresponding sum of mapped reads for a given library and multiplying by 20,000,000. In *dcl5* libraries, called 24-*PHAS* loci were compared with reference loci that retained detectable abundance but did not pass the high-confidence 24-*PHAS* calling criteria.

Candidate loci were also visualized in *Phasis*-score and Howell-score space. Validated 24-*PHAS* calls, filtered false-positive candidates, *dcl5*-associated candidates, and non-*PHAS* background loci were plotted relative to the fixed-threshold criteria used for 24-*PHAS* calling. This diagnostic was used to evaluate whether candidate loci showed concordant statistical and register-level support or instead represented threshold-conflict candidates with incomplete evidence for high-confidence 24-*PHAS* annotation. CP20M-normalized abundance distributions were summarized descriptively using counts and median CP20M values for called 24-*PHAS* loci and reference-overlapping non-called loci.

### Additional Plant Validation Datasets

For petunia and columbine, individual-library and pooled-library *Phasis* outputs were compared with the corresponding curated reproductive 24-*PHAS* reference sets using *PhasMatch*. Performance was summarized using call-level precision, reference-locus recall, F1 score, and reference-side overlap between evidence modes.

For Arabidopsis, individual-library and pooled-library *Phasis* outputs were compared with Araport11 *TAS* annotations to evaluate recovery of annotated *TAS*-associated 21-*PHAS* loci. Pooled Arabidopsis 24-*PHAS* candidates were manually inspected against Araport11 gene and transposable-element annotations. Candidates that lacked convincing phased-register structure or overlapped annotated transposable-element genes were classified as unsupported rather than as bona fide reproductive 24-*PHAS* loci.

### Drosophila Phase-Length Scan and Annotation Audit

Drosophila ovarian sRNA libraries were analyzed as a non-plant boundary test because animal piRNAs can show phased production through Zucchini-dependent processing, but this pathway differs mechanistically from plant DCL-dependent phasiRNA biogenesis. The analysis used ovarian libraries from Han et al. (2015) and the corresponding FlyBase release 6.54 genome assembly. Because piRNAs have a broader size distribution than plant phasiRNAs, Phasis was run across phase lengths from 23 to 30 nt.

Automated calls from the Drosophila phase-length scan were manually inspected to determine whether they showed plant-like *PHAS* architecture or instead reflected local abundance artifacts, repeat-associated sRNAs, or piRNA-sized signals that could mimic fixed-register periodicity. For the three candidates reported by *Phasis*, locus plots were examined for coherent locus-level organization.

Candidate coordinates were then compared with FlyBase release 6.54 gene, transposable-element, repeat, and other genomic annotations. Calls were interpreted as unsupported when their apparent periodicity could be explained by repeat-rich or transposable-element-associated sequence context, concentrated local abundance, piRNA-associated sequence features such as 1U status where relevant, or other patterns inconsistent with plant-like *PHAS* architecture.

### Comparison with Existing Tools in Matched Pooled-Library Analyses

To evaluate *Phasis* relative to existing *PHAS*-discovery tools, we performed matched pooled-library benchmarks using *Phasis*, *ShortStack* v3.8.5, and *PhaseTank* v1.0 outputs generated from the same sRNA-library sets and evaluated against the same curated reference sets. Direct tool-to-tool comparisons were restricted to pooled-library analyses because the comparator tools do not provide the same individual-library evidence mode as *Phasis*. Individual-library *Phasis* results were therefore not used for direct comparator benchmarking.

For each species and phase class, *PHAS* calls from each tool were compared with the corresponding curated reference set using *PhasMatch*. Benchmarking was summarized using call-level and reference-locus metrics. Using the metric definitions described above, call-level precision quantified the proportion of calls supported by curated reference loci, whereas reference-locus recall quantified recovery of curated loci. F1 score summarized the balance between call-level precision and reference-locus recall. For *ShortStack*, positive calls were restricted to records with PhaseScore ≥ 12.5, and for *PhaseTank*, positive calls were restricted to records with Phased_Score ≥ 12.5. Although these metrics are named differently across software packages, all are derived from the phased-locus scoring framework originally described by Howell et al. (2007). Each tool implements its own adaptation of that framework, resulting in software-specific score calculations and nomenclature. These threshold values were selected to match the Howell-score cutoff used for *Phasis* positive-call filtering before benchmarking. The main comparator benchmark table is provided as Supplemental Table S1.

## Acknowledgments

We thank Chong Teng, Jianing Wang, and Jo-Wei Hsieh for their testing of *Phasis* and for providing valuable suggestions that helped improve the software’s functionality and usability. We are also grateful to Mayumi Nakano for her earlier work in developing a phasiRNA visualization tool, which provided useful ideas and code that informed parts of the current implementation.

## Data Availability

The sequencing datasets analyzed in this study are available from Gene Expression Omnibus (GEO) or the Sequence Read Archive (SRA). Benchmark summary tables, sequencing-library metadata and accession identifiers, curated *PHAS* locus annotations and coordinates, and representative analysis workflows are provided in Supplemental Tables S1–S5 and Supplemental File S1.

## Code Availability

*Phasis* is an open-source Python software package available from the project GitHub repository https://github.com/blakemeyerslab/Phasis under the OSI Artistic License 2.0. The archived, versioned release corresponding to the implementation used in this study is available at Zenodo (DOI: 10.5281/zenodo.20435283). The software was tested with Python 3.12 on Unix-like operating systems, including Linux and macOS. Installation is provided via pip from the repository, which installs the *Phasis* command-line interface; *HISAT2* and *SAMtools* must be available on the user’s PATH. A detailed user manual, including installation instructions, system requirements, command-line options, and example analysis workflows, is provided in the repository.

## Supplemental Material

Supplemental Material, including Supplemental Methods, Supplemental Fig. S1, Supplemental Tables S1–S5, and Supplemental File S1, is available for this article.

## Author Contributions Statement

AK conceived the project. THCR, AK, BCM, SL, and VM designed and implemented the software. THCR, AK, and SL performed benchmarking analyses, curated reference comparisons, and evaluated validation datasets. THCR, AK, SL, and BCM interpreted results and wrote the manuscript. All authors read and approved the final manuscript.

## Supplemental Methods

### Overview of the Phasis Analysis Framework

Phasis is an open-source software tool for de novo discovery, scoring, classification, and interpretation of phased small RNA (*PHAS*) loci from plant small RNA sequencing data. The implementation used in this study analyzes small RNA libraries together with a reference genome or transcriptome and reports candidate *PHAS* loci, locus-level evidence metrics, diagnostic visualizations, and phased small RNA unit annotations. The workflow is organized into two major analytical stages. First, Phasis performs Initial *PHAS* Candidate Detection using an established cluster-level framework. This stage maps small RNA libraries, identifies small RNA-producing clusters, evaluates locus-level features associated with phased biogenesis, computes phasing-related evidence metrics, and applies GMM-supported candidate detection with configurable feature thresholds. Second, candidate loci are evaluated by a Register-Resolved Locus Interpretation Layer. This post-detection layer examines candidate loci at single-register resolution and reconstructs the organization of phased signal within each locus. It interprets and stratifies candidate loci after detection by identifying the main phased unit, register anchor, coherent same-register extensions, accepted opposite-strand partners, secondary phased windows, overlapping alternative windows, and *PHAS*-like evidence categories. The main manuscript summarizes the conceptual workflow and key benchmark outcomes, whereas the sections below provide implementation-oriented details.

### Input Small RNA Libraries and Reference Sequences

Phasis accepts user-supplied small RNA libraries together with a reference genome or transcriptome. Small RNA libraries can be provided in common sequence formats (FASTA and FASTQ) or as tag-count files containing deduplicated small RNA sequences and their abundances. Tag-count input stores each unique small RNA sequence once together with its count and is useful for large datasets.

For the analyses reported in this study, small RNA libraries were processed with the reference assemblies and sample groupings used for each validation or benchmarking dataset. These included maize, coffee, *Arabidopsis*, petunia, columbine, and *Drosophila* datasets described in the main manuscript. Dataset source information and accession identifiers are summarized in the main manuscript and Supplemental Table S4.

### Read Mapping and Small RNA Cluster Construction

Small RNA reads are aligned to the selected reference using HISAT2 (Kim et al. 2019) with settings appropriate for short, unspliced small RNA alignments. Alignments are generated with no soft clipping and no spliced alignment and sorted with SAMtools (Li et al. 2009) before downstream cluster discovery.

After mapping, Phasis identifies genomic or transcript-derived regions enriched for small RNAs of the expected phase length, typically 21 nt or 24 nt in plant datasets. Adjacent small RNA-producing regions are merged or separated according to user-configurable cluster parameters. These candidate small RNA clusters form the basic locus units for downstream feature calculation, phasing-score evaluation, candidate classification, and register-resolved interpretation.

Important user-configurable parameters include the expected phasing periodicity, the minimum read depth required for cluster consideration, the buffer distance used to separate adjacent clusters, and the maximum number of permitted reference alignments per small RNA read. In the analyses reported here, parameter settings were selected according to the intended phase class and dataset context. Exact command examples are provided in Supplemental File S1.

### Cluster-Level Features Used for Initial PHAS Candidate Detection

For each candidate small RNA cluster, Phasis calculates locus-level features expected to distinguish *PHAS* loci from other small RNA-producing regions. These features include strand bias, cluster complexity, abundance normalized by locus length, the proportion of reads matching the expected phase length, the Howell score, and the Phasis score. Here, we use Howell score to refer to the adapted phased-score metric originally described by Howell et al. (2007). In Phasis, this score is modified to allow strand-specific positional variance of plus or minus one nucleotide at the 5′ end of the expected phase register, reflecting limited positional variation that can occur during DCL-mediated processing. Strand bias summarizes the distribution of mapped small RNA abundance between genomic strands. Cluster complexity summarizes the diversity of distinct small RNA fragments within a cluster relative to total abundance. Length-normalized abundance provides a measure of small RNA accumulation adjusted for locus length. The phase-length ratio captures the fraction of reads corresponding to the expected phased small RNA size class. Together, these features represent each cluster in a shared evidence space used for initial candidate detection, GMM-supported stratification, threshold filtering, and downstream register-resolved interpretation.

### Phasis Score Calculation

The Phasis score quantifies statistical evidence for periodic small RNA accumulation within a candidate cluster. For each cluster, Phasis evaluates candidate phased registers using a sliding-window strategy across both strands and across the expected phasing periodicity.

Within each evaluated window, Phasis applies two complementary statistical tests. A hypergeometric test evaluates enrichment of small RNA reads at expected phased positions relative to non-phased positions. A Wilcoxon rank-sum test compares read abundance at expected phased and non-phased positions. These tests capture complementary aspects of phasing: positional enrichment and abundance structure.

P-values from these two tests are combined within evaluated windows using Stouffer’s method. Window-level evidence is then aggregated at the candidate-locus level using Fisher’s method. The resulting combined p-value is transformed as −log10(p) to produce the Phasis score, with higher values indicating stronger statistical support for phased small RNA accumulation. Because multi-window p-value aggregation can produce combined p-values near the lower numerical range of floating-point representation, Phasis reports scores on a capped scale, with values ≥300 recorded as 300. This cap prevents underflow- or precision-related artifacts in the extreme tail while preserving the distinction between weak, moderate, and strong phasing support. Statistical calculations are implemented using SciPy (Virtanen et al. 2020).

### GMM-Supported Initial Candidate Detection

Initial *PHAS* Candidate Detection uses the cluster-level evidence framework to distinguish candidate *PHAS* loci from other sRNA-producing clusters before register-resolved interpretation. In the implementation used for this study, Phasis applies an unsupervised Gaussian mixture model (GMM) to locus-level evidence features, including the Phasis score, strand bias, cluster complexity, length-normalized abundance, and the phase-length abundance ratio. The GMM provides an initial data-driven stratification of candidate clusters but is not used as the sole determinant of final high-confidence *PHAS* calls. Classifier implementation uses scikit-learn (Pedregosa et al. 2011).

After GMM-supported initial detection, Phasis applies configurable feature thresholds and passes retained candidates to the Register-Resolved Locus Interpretation Layer. In the default configuration evaluated in this study, final high-confidence *PHAS* calls therefore depend on the combination of locus-level evidence, threshold filtering, and register-resolved architecture. This design allows the initial detection step to remain broad while RRL evaluates whether candidate loci contain coherent phased structure consistent with plant DCL-dependent phasiRNA biogenesis.

### Register-Resolved Locus Interpretation Layer

After Initial *PHAS* Candidate Detection, candidate loci are evaluated by the Register-Resolved Locus Interpretation Layer (RRL). This layer provides post-detection interpretation of local phasing architecture within each candidate locus. It examines the local register-level Howell-score trace. The highest Howell-score-supported register is designated as the register anchor, also referred to as the HPSP. This anchor defines the primary phased register for the candidate locus. Phasis then quantifies exact-register support around the anchor and evaluates whether additional same-register phased signal extends coherently from the anchored region.

The anchored structure defines the main phased unit, which is the primary register-resolved phased window associated with the candidate locus. Additional same-register phased signal connected to this unit is reported as a coherent extension. Phasis also tests for an accepted main opposite-strand partner, defined as an opposite-strand phased signal compatible with canonical phased duplex geometry. Opposite-strand partner evaluation is based on register geometry and local phased evidence. It is not intended to infer transcript strand or trigger-miRNA strand.

Candidate *PHAS* loci may contain additional supported phased signals beyond the main phased unit. When such signals occur outside the main unit, they can be promoted as secondary phased windows. When additional supported signal overlaps the main locus but is supported by an alternative register or shifted phased structure, it can be reported as an overlapping alternative window. These categories preserve local phased architecture without assuming that every promoted signal is an independent biological locus.

### PHAS-Like Evidence Category

The Register-Resolved Locus Interpretation Layer supports an intermediate *PHAS*-like evidence category. *PHAS*-like loci are candidates with detectable phased structure that do not meet the register-level evidence required for a final high-confidence *PHAS* call.

This category is intended to preserve potentially informative lower-confidence phased structures rather than discarding them as failed candidates. *PHAS*-like classification can arise when exact-register support is weak, when the local phasing context is crowded, when alternative registers compete with the main phased unit, or when evidence is detectable but insufficiently coherent for a final *PHAS* interpretation. In the implementation used here, candidates with no exact-register support at the register anchor were not promoted to high-confidence *PHAS* calls, even when relaxed ±1-nt support was detected, because relaxed support alone was considered insufficient to define a coherent primary phased register. In downstream analyses, *PHAS*-like loci should therefore be treated separately from high-confidence *PHAS* calls.

### Diagnostic Plots and Phased-Unit Annotation Tables

Register-resolved interpretation supports locus-specific diagnostic plots and phased small RNA unit annotations. These outputs are designed to report not only whether a locus is classified as *PHAS*, but also how phased signal is organized within the locus.

The diagnostic plots separate abundance context from score context. They display the main phased unit, highlight accepted opposite-strand partners, and show promoted secondary or overlapping alternative phased windows when present. These plots are interpretive diagnostics for candidate loci and do not by themselves change the initial candidate-detection, threshold-filtering, or register-resolved interpretation rules.

The corresponding phased small RNA tables record unit membership and role for phased small RNAs assigned to an interpreted locus. These annotations allow downstream analyses to distinguish small RNAs assigned to the main phased unit, coherent extension, opposite-strand partner, secondary phased window, or overlapping alternative window. This register-resolved annotation provides a structured representation of locus architecture for downstream summary, visualization, and biological interpretation.

### Benchmarking and Curated-Reference Evaluation

Benchmarking analyses compare Phasis calls against curated reference sets for each species and phase class. Candidate loci are matched to reference loci using coordinate-overlap criteria. True positives, false positives, and false negatives are then used to calculate precision, recall, and F1 score. Reference-set curation and false-positive review included manual inspection of phased small RNA evidence using locus-level diagnostic plots and the MPSS phasing analysis browser (Nakano et al. 2020). This inspection was used to confirm that curated reference loci showed convincing phased evidence and to evaluate whether candidate calls from Phasis or comparator tools represented coherent phased loci rather than unsupported sRNA-producing regions.

For comparative benchmarking, Phasis and external tools were evaluated using the same reference assemblies, input datasets, curated reference sets, and matching criteria whenever possible. PhaseTank v1.0 and ShortStack v3.8.5 were used as comparators because they are commonly used tools that report phased small RNA loci or phased-locus scores and therefore provide directly benchmarkable outputs for *PHAS*-locus calling (Guo et al. 2015; Axtell 2013). This design was intended to ensure that differences in performance reflected differences in locus calling and scoring behavior rather than differences in input data, reference sets, or evaluation criteria.

### Dataset-Specific Benchmarking Contexts

Maize and coffee 21-*PHAS* analyses use curated loci and matched non-*PHAS* clusters derived from datasets with clear library-specific phasing signals (Zhai et al. 2015; Cherubino Ribeiro et al. 2024). These datasets support the maize and coffee benchmark analyses and the cross-species validation reported in this study.

Maize 24-*PHAS* analyses encompass W23 2-mm anther data and *dcl5* mutant material as a negative reference for DCL5-dependent 24-nt reproductive phasiRNAs. The biology of DCL5-dependent 24-nt reproductive phasiRNAs and the potential for false-positive 24-nt siRNA-dominated *PHAS* annotations have been previously described (Polydore et al. 2018; Zhai et al. 2015; Liu et al. 2020).

Petunia and columbine 24-*PHAS* analyses used bud or reproductive datasets and reference genomes described in previous studies (Bombarely et al. 2016; Filiault et al. 2018; Xia et al. 2019). These datasets tested whether Phasis could recover reproductive 24-*PHAS* loci in additional angiosperm systems beyond the maize and coffee benchmark contexts, and whether the default evidence thresholds and register-resolved interpretation remained effective across distinct reproductive 24-*PHAS* genomic contexts.

Arabidopsis analyses use seedling libraries to evaluate recovery of known 21-*PHAS*/*TAS* loci and the expected absence of miR2275-dependent 24-nt reproductive *PHAS* signals (Li et al. 2016; Reiser et al. 2024; Xia et al. 2019). Drosophila ovarian small RNA libraries are used as a scope-boundary test because animal piRNA phasing is mechanistically distinct from plant DCL-dependent phasiRNA biogenesis (Han et al. 2015; Mohn et al. 2015; Öztürk-Çolak et al. 2024).

### Statistical Analyses and Performance Metrics

Statistical analyses used by Phasis are implemented with standard scientific Python libraries. Phasing evidence is evaluated using the hypergeometric and Wilcoxon rank-sum tests described above, with p-value combination by Stouffer’s and Fisher’s methods. Dataset-level benchmarking is reported using call-level precision, reference-locus recall, and F1 score derived from true-positive calls, false-positive calls, and unrecovered reference loci. For benchmarking against curated loci, each candidate is categorized as true-positive or false-positive based on the defined coordinate-overlap criteria, and each reference locus is categorized as recovered or unrecovered according to whether it is overlapped by one or more calls.

### Reproducibility and Archived Software Release

The software version, command-line settings, reference files, input libraries, and output summaries used for the final analyses are provided through the project repository, the archived software release, Supplemental File S1, and Supplemental Tables S1–S5.

---

**Figure S1.**
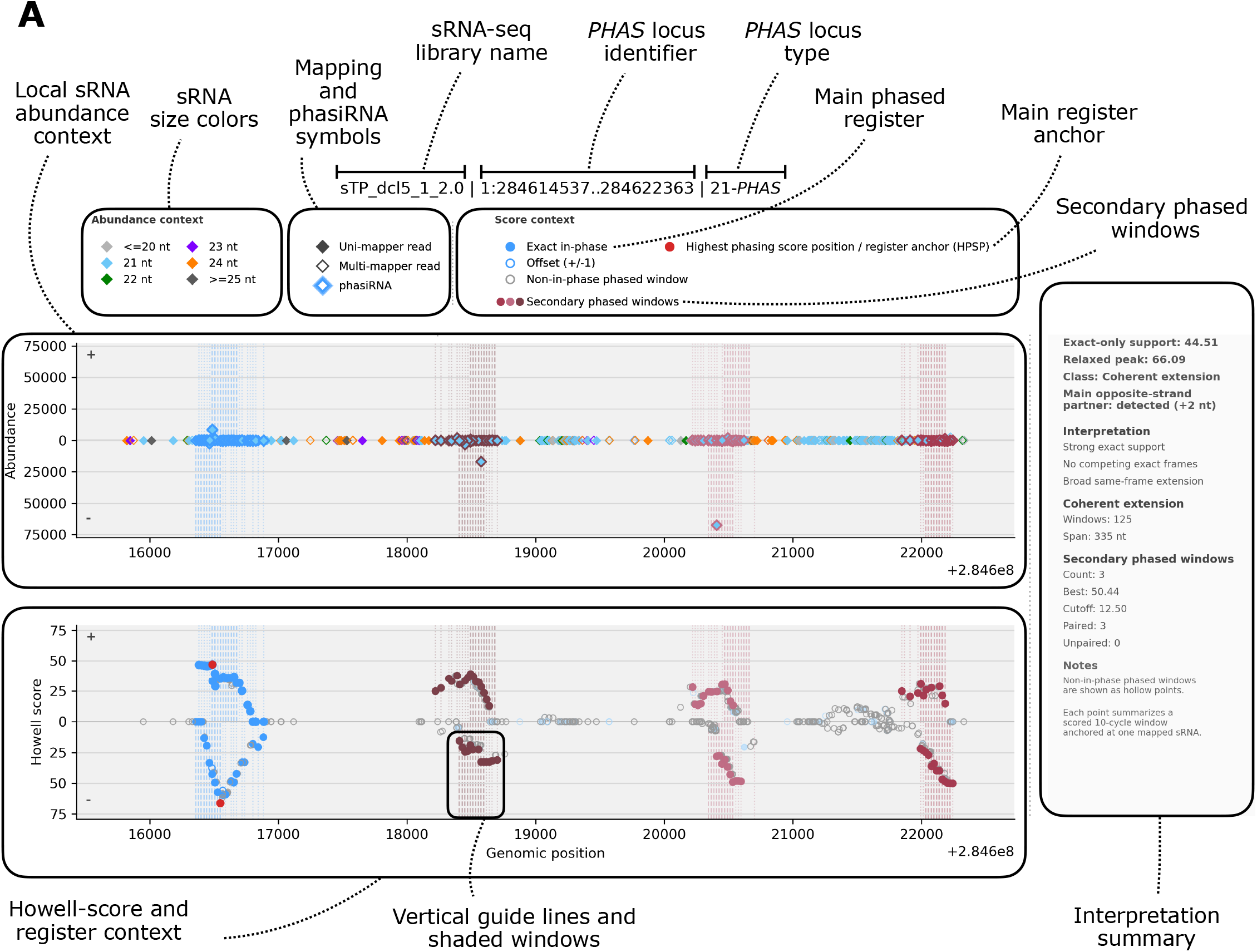
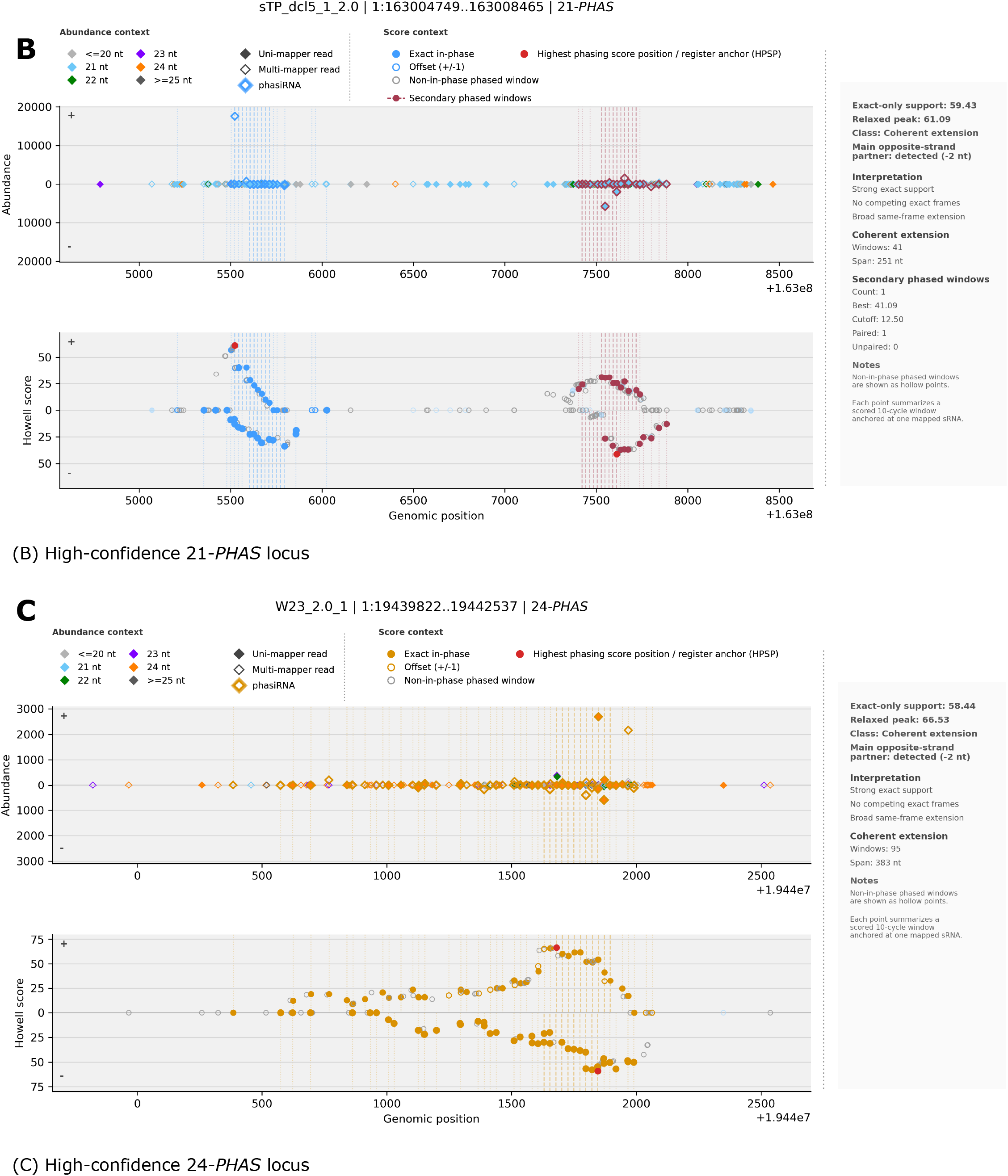
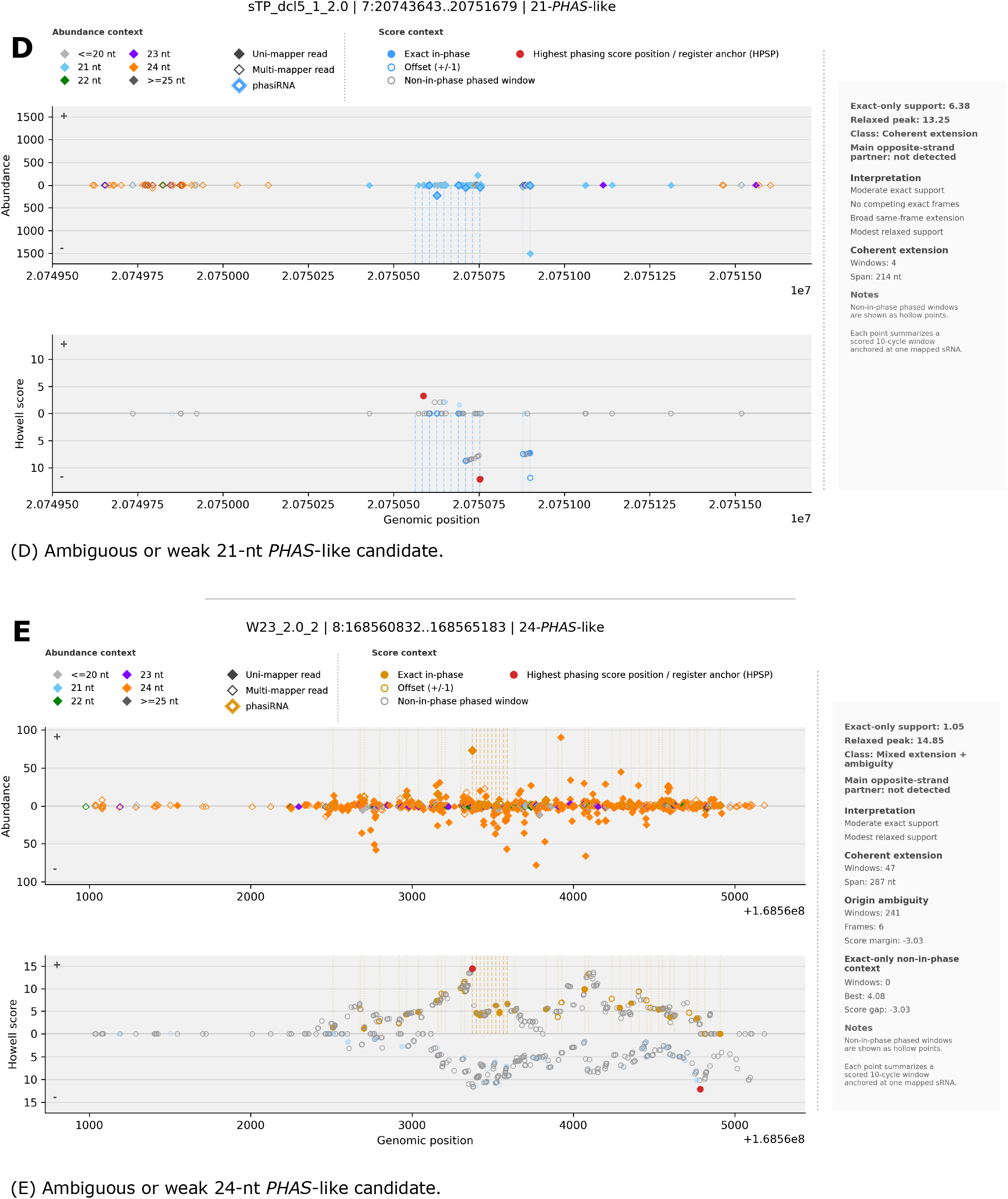
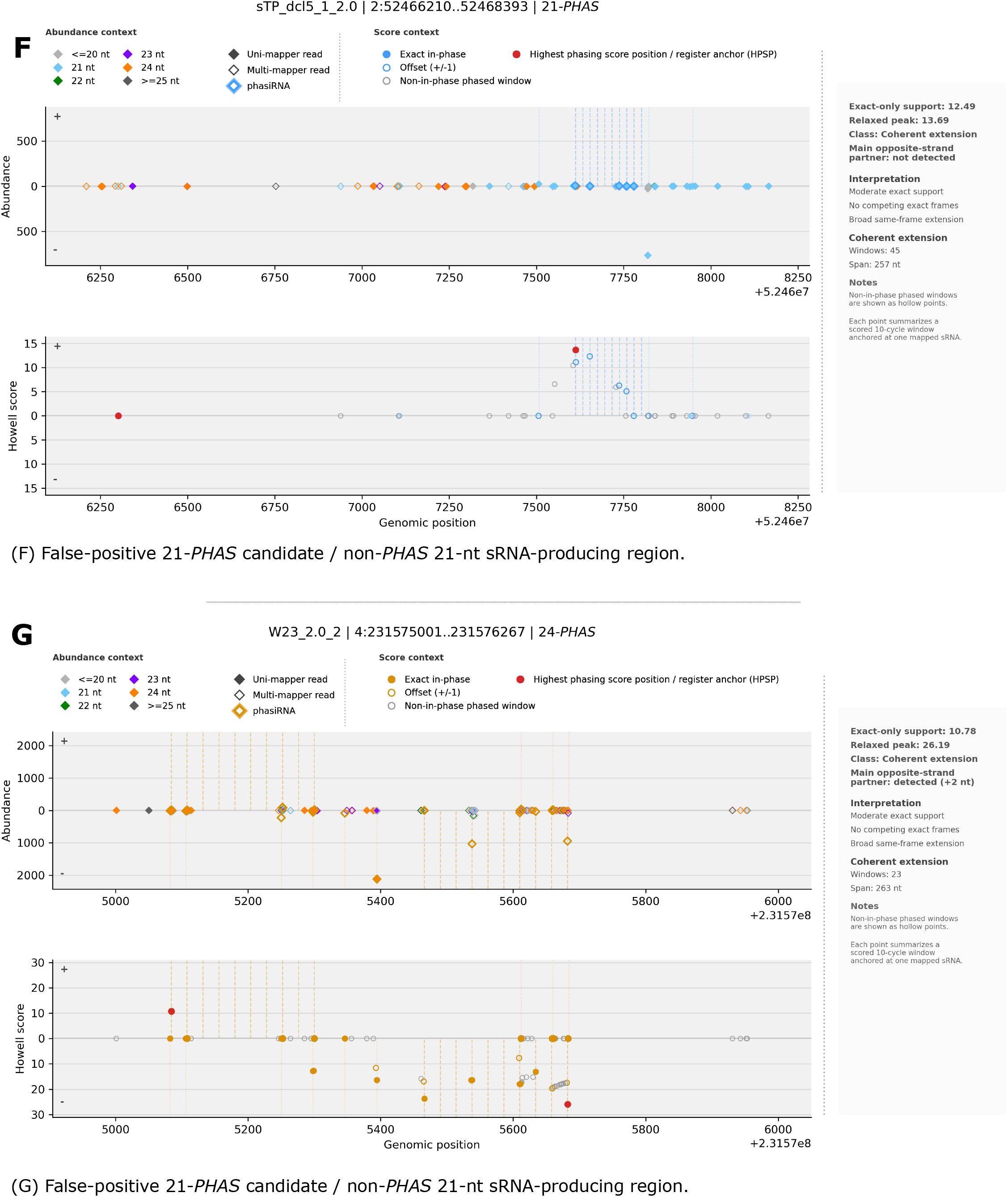
Guide and representative examples of *Phasis* locus-level diagnostic plots. Guide to *Phasis* locus-level diagnostic plots. The title line identifies the sRNA-seq library, genomic interval, and predicted *PHAS* locus type. The top panel shows local sRNA abundance context: each diamond represents a mapped sRNA at its genomic position, with reads on opposite strands plotted above and below the horizontal axis. Diamond color indicates sRNA size class, allowing visual assessment of whether the locus is dominated by the expected 21- or 24-nt species. Filled diamonds indicate uniquely mapped reads, open diamonds indicate multi-mapped reads, and larger outlined diamonds indicate phasiRNA reads assigned by *Phasis* to the called phased register. Genomic coordinates are shown on the x-axis, with the full chromosome interval provided in the plot title; when an axis offset is displayed at the lower right, tick labels are interpreted relative to that offset. The bottom panel shows Howell-score and register context. Each point represents a scored phased window anchored at a mapped sRNA position, with the y-axis showing Howell-score support on each strand. Filled colored points indicate exact in-phase support for the main phased register, open colored points indicate windows offset by ±1 nt from that register, and grey open points indicate phased windows not assigned to the main register. The red point marks the highest phasing-score position, or main register anchor. Additional supported phased windows are shown in secondary colors; these colors distinguish secondary phased-window groups and do not indicate sRNA size. Vertical guide lines and shaded windows connect abundance patterns in the top panel with the scored register evidence in the bottom panel. The right-hand summary box reports locus-level support, RRL interpretation, coherent-extension information, secondary-window statistics, and notes used for manual review. (A) Annotated guide to the plot elements used in *Phasis* diagnostic visualizations. The top panel shows local sRNA abundance, strand orientation, size class, mapping status, and reads assigned to the called phased register. The bottom panel shows Howell-score and register support across scored phased windows, including the main phased register, ±1-nt offset windows, non-main phased windows, the highest phasing-score position/register anchor, and secondary phased windows. The right-hand box summarizes locus-level evidence and RRL interpretation used for manual review.

## References

2. Allen E, Xie Z, Gustafson AM, Carrington JC. 2005. microRNA-Directed Phasing during *Trans*-Acting siRNA Biogenesis in Plants. Cell 121: 207–221.

3. Anleu Gil MX, Meyers BC. 2026. Reproductive phasiRNAs are the piRNAs of plants. Trends Genet 42: 268–281.

4. Axtell MJ. 2013. ShortStack: Comprehensive annotation and quantification of small RNA genes. RNA 19: 740–751.

5. Bombarely A, Moser M, Amrad A, Bao M, Bapaume L, Barry CS, Bliek M, Boersma MR, Borghi L, Bruggmann R, et al. 2016. Insight into the evolution of the Solanaceae from the parental genomes of *Petunia hybrida*. Nat Plants 2: 16074.

6. Bu Y, Zheng J, Jia C. 2023. An efficient deep learning based predictor for identifying miRNA-triggered phasiRNA loci in plant. Math Biosci Eng 20: 6853–6865.

7. Cherubino Ribeiro TH, Baldrich P, De Oliveira RR, Fernandes-Brum CN, Mathioni SM, De Sousa Cardoso TC, De Souza Gomes M, Do Amaral LR, Pimenta De Oliveira KK, Dos Reis GL, et al. 2024. The floral development of the allotetraploid *Coffea arabica* L. correlates with a small RNA dynamic reprogramming. Plant J 118: 1848–1863.

8. Fei Q, Xia R, Meyers BC. 2013. Phased, secondary, small interfering RNAs in posttranscriptional regulatory networks. Plant Cell 25: 2400–2415.

9. Feng Z, Feng J, Zhang B, Fei Y, Zhang H, Huang J. 2023. *PhasiHunter*: a robust phased siRNA regulatory cascade mining tool based on multiple reference sequences. Bioinformatics 39: btad676.

10. Filiault DL, Ballerini ES, Mandáková T, Aköz G, Derieg NJ, Schmutz J, Jenkins J, Grimwood J, Shu S, Hayes RD, et al. 2018. The *Aquilegia* genome provides insight into adaptive radiation and reveals an extraordinarily polymorphic chromosome with a unique history. eLife 7: e36426.

11. Guo Q, Qu X, Jin W. 2015. *PhaseTank*: Genome-wide computational identification of phasiRNAs and their regulatory cascades. Bioinformatics 31: 284–286.

12. Han BW, Wang W, Li C, Weng Z, Zamore PD. 2015. piRNA-guided transposon cleavage initiates Zucchini-dependent, phased piRNA production. Science 348: 817–821.

13. Howell MD, Fahlgren N, Chapman EJ, Cumbie JS, Sullivan CM, Givan SA, Kasschau KD, Carrington JC. 2007. Genome-Wide Analysis of the RNA-DEPENDENT RNA POLYMERASE6/DICER-LIKE4 Pathway in *Arabidopsis* Reveals Dependency on miRNA- and tasiRNA-Directed Targeting. Plant Cell 19: 926–942.

14. Johnson C, Kasprzewska A, Tennessen K, Fernandes J, Nan G-L, Walbot V, Sundaresan V, Vance V, Bowman LH. 2009. Clusters and superclusters of phased small RNAs in the developing inflorescence of rice. Genome Res 19: 1429–1440.

15. Kakrana A, Li P, Patel P, Hammond R, Anand D, Mathioni SM, Meyers BC. 2017. *PHASIS*: A computational suite for de novo discovery and characterization of phased, siRNA-generating loci and their miRNA triggers. *bioRxiv* 158832.

16. Li S, Le B, Ma X, Li S, You C, Yu Y, Zhang B, Liu L, Gao L, Shi T. 2016. Biogenesis of phased siRNAs on membrane-bound polysomes in *Arabidopsis*. eLife 5: e22750.

17. Marin E, Jouannet V, Herz A, Lokerse AS, Weijers D, Vaucheret H, Nussaume L, Crespi MD, Maizel A. 2010. miR390, *Arabidopsis TAS3* tasiRNAs, and Their *AUXIN RESPONSE FACTOR* Targets Define an Autoregulatory Network Quantitatively Regulating Lateral Root Growth. Plant Cell 22: 1104–1117.

18. Mohn F, Handler D, Brennecke J. 2015. piRNA-guided slicing specifies transcripts for Zucchini-dependent, phased piRNA biogenesis. Science 348: 812–817.

19. Öztürk-Çolak A, Marygold SJ, Antonazzo G, Attrill H, Goutte-Gattat D, Jenkins VK, Matthews BB, Millburn G, dos Santos G, Tabone CJ, et al. 2024. FlyBase: updates to the *Drosophila* genes and genomes database. Genetics 227: iyad211.

20. Peragine A, Yoshikawa M, Wu G, Albrecht HL, Poethig RS. 2004. *SGS3* and *SGS2/SDE1/RDR6* are required for juvenile development and the production of trans-acting siRNAs in *Arabidopsis*. Genes Dev 18: 2368–2379.

21. Polydore S, Lunardon A, Axtell MJ. 2018. Several phased siRNA annotation methods can frequently misidentify 24 nucleotide siRNA-dominated PHAS loci. Plant Direct 2: e00101.

22. Seong K, Kumar R, Prigozhin DM, Lunde C, Cherubino Ribeiro TH, Bélanger S, Hsieh J-WA, Tang M, Meyers BC, Krasileva KV. 2026. The annotated blueprint: integrated functional genomic resources for a model tetraploid wheat *Triticum turgidum* cv. Kronos. New Phytol. 10.1111/nph.71006.

23. Teng C, Zhang H, Hammond R, Huang K, Meyers BC, Walbot V. 2020. Dicer-like 5 deficiency confers temperature-sensitive male sterility in maize. Nat Commun 11: 2912.

24. Vazquez F, Vaucheret H, Rajagopalan R, Lepers C, Gasciolli V, Mallory AC, Hilbert J-L, Bartel DP, Crété P. 2004. Endogenous *trans*-acting siRNAs regulate the accumulation of *Arabidopsis* mRNAs. Mol Cell 16: 69–79.

25. Xia R, Chen C, Pokhrel S, Ma W, Huang K, Patel P, Wang F, Xu J, Liu Z, Li J, et al. 2019a. 24-nt reproductive phasiRNAs are broadly present in angiosperms. Nat Commun 10: 627.

26. Zhai J, Jeong D-H, De Paoli E, Park S, Rosen BD, Li Y, González AJ, Yan Z, Kitto SL, Grusak MA. 2011. MicroRNAs as master regulators of the plant *NB-LRR* defense gene family via the production of phased, *trans*-acting siRNAs. Genes Dev 25: 2540–2553.

27. Zhai J, Zhang H, Arikit S, Huang K, Nan GL, Walbot V, Meyers BC. 2015. Spatiotemporally dynamic, cell-type-dependent premeiotic and meiotic phasiRNAs in maize anthers. Proc Natl Acad Sci U S A 112: 3146–3151.

28. Zhan J, Meyers BC. 2023. Plant Small RNAs: Their Biogenesis, Regulatory Roles, and Functions. Annu Rev Plant Biol 74: 21–51.

## References for Supplemental Methods

29. Axtell MJ. 2013. *ShortStack*: Comprehensive annotation and quantification of small RNA genes. RNA 19: 740–751.

30. Bombarely A, Moser M, Amrad A, Bao M, Bapaume L, Barry CS, Bliek M, Boersma MR, Borghi L, Bruggmann R, et al. 2016. Insight into the evolution of the Solanaceae from the parental genomes of *Petunia hybrida*. Nat Plants 2: 16074.

31. Cherubino Ribeiro TH, Baldrich P, De Oliveira RR, Fernandes-Brum CN, Mathioni SM, De Sousa Cardoso TC, De Souza Gomes M, Do Amaral LR, Pimenta De Oliveira KK, Dos Reis GL, et al. 2024. The floral development of the allotetraploid *Coffea arabica* L. correlates with a small RNA dynamic reprogramming. Plant J 118: 1848–1863.

32. Filiault DL, Ballerini ES, Mandáková T, Aköz G, Derieg NJ, Schmutz J, Jenkins J, Grimwood J, Shu S, Hayes RD, et al. 2018. The *Aquilegia* genome provides insight into adaptive radiation and reveals an extraordinarily polymorphic chromosome with a unique history. eLife 7: e36426.

33. Guo Q, Qu X, Jin W. 2015. *PhaseTank*: Genome-wide computational identification of phasiRNAs and their regulatory cascades. Bioinformatics 31: 284–286.

34. Han BW, Wang W, Li C, Weng Z, Zamore PD. 2015. piRNA-guided transposon cleavage initiates Zucchini-dependent, phased piRNA production. Science 348: 817–821.

35. Howell MD, Fahlgren N, Chapman EJ, Cumbie JS, Sullivan CM, Givan SA, Kasschau KD, Carrington JC. 2007. Genome-Wide Analysis of the RNA-DEPENDENT RNA POLYMERASE6/DICER-LIKE4 Pathway in *Arabidopsis* Reveals Dependency on miRNA-and tasiRNA-Directed Targeting. Plant Cell 19: 926–942.

36. Kim D, Paggi JM, Park C, Bennett C, Salzberg SL. 2019. Graph-based genome alignment and genotyping with *HISAT2* and *HISAT*-genotype. Nat Biotechnol 37: 907–915.

37. Li H, Handsaker B, Wysoker A, Fennell T, Ruan J, Homer N, Marth G, Abecasis G, Durbin R. 2009. The sequence alignment/map format and *SAMtools*. Bioinformatics 25: 2078–2079.

38. Li S, Le B, Ma X, Li S, You C, Yu Y, Zhang B, Liu L, Gao L, Shi T. 2016. Biogenesis of phased siRNAs on membrane-bound polysomes in *Arabidopsis*. eLife 5: e22750.

39. Liu Y, Teng C, Xia R, Meyers BC. 2020. PhasiRNAs in plants: their biogenesis, genic sources, and roles in stress responses, development, and reproduction. Plant Cell 32: 3059–3080.

40. Mohn F, Handler D, Brennecke J. 2015. piRNA-guided slicing specifies transcripts for Zucchini-dependent, phased piRNA biogenesis. Science 348: 812–817.

41. Nakano M, McCormick K, Demirci C, Demirci F, Gurazada SGR, Ramachandruni D, Dusia A, Rothhaupt JA, Meyers BC. 2020. Next-generation sequence databases: RNA and genomic informatics resources for plants. Plant Physiol 182: 136–146.

42. Öztürk-Çolak A, Marygold SJ, Antonazzo G, Attrill H, Goutte-Gattat D, Jenkins VK, Matthews BB, Millburn G, dos Santos G, Tabone CJ, et al. 2024. FlyBase: updates to the *Drosophila* genes and genomes database. Genetics 227: iyad211.

43. Pedregosa F, Varoquaux G, Gramfort A, Michel V, Thirion B, Grisel O, Blondel M, Prettenhofer P, Weiss R, Dubourg V, et al. 2011. Scikit-learn: Machine Learning in Python. J Mach Learn Res 12: 2825–2830.

44. Polydore S, Lunardon A, Axtell MJ. 2018. Several phased siRNA annotation methods can frequently misidentify 24 nucleotide siRNA-dominated *PHAS* loci. Plant Direct 2: e00101.

45. Reiser L, Bakker E, Subramaniam S, Chen X, Sawant S, Khosa K, Prithvi T, Berardini TZ. 2024. The *Arabidopsis* Information Resource in 2024. Genetics 227: iyae027.

46. Virtanen P, Gommers R, Oliphant TE, Haberland M, Reddy T, Cournapeau D, Burovski E, Peterson P, Weckesser W, Bright J, et al. 2020. SciPy 1.0: fundamental algorithms for scientific computing in Python. Nat Methods 17: 261–272.

47. Xia R, Chen C, Pokhrel S, Ma W, Huang K, Patel P, Wang F, Xu J, Liu Z, Li J, et al. 2019. 24-nt reproductive phasiRNAs are broadly present in angiosperms. Nat Commun 10: 627.

48. Zhai J, Zhang H, Arikit S, Huang K, Nan GL, Walbot V, Meyers BC. 2015. Spatiotemporally dynamic, cell-type-dependent premeiotic and meiotic phasiRNAs in maize anthers. Proc Natl Acad Sci U S A 112: 3146–3151.

